# Personal network features are embodied in real-time social interaction dynamics

**DOI:** 10.64898/2026.07.03.736342

**Authors:** Aliaksandr Dabranau, Sune Lehmann, Ivana Konvalinka

## Abstract

People navigate dozens of social interactions daily. Through these, they construct and maintain diverse social networks. While personal dispositions drive how people interact and connect others, the resulting networks feed back into daily social experiences by creating interaction opportunities and modulating cognitive processes. Here, we show that people’s moment-to-moment interaction behaviors are informative of the structure of their personal social network, even if the interaction occurs between strangers. We paired 104 strangers to perform an interactive motor task and investigated whether emergent leader-follower roles and spontaneous movement synchronization reflected similarities or differences in their personal networks. Participants with tighter networks (smaller, denser, more constrained, and with fewer communities) relative to the interaction partner adapted their movements to the partner more. Using multivariate temporal response functions (mTRF) to measure neural encoding of self and other’s movements, we found that, conversely, participants with less tight networks exhibited higher neural encoding of their partner’s movements. This suggests that the network effect on behavioral adaptation was unlikely to be driven by heightened monitoring of the partner. Higher spontaneous movement synchronization was associated with higher personality agreeableness and personality similarity between interacting partners. Our results demonstrate that signatures of personal social environment manifest dynamically in real-time behaviors and that these effects are independent of personality effects. Thus, social networks set an additional context for interpersonal interaction even for partners without a prior relationship.

## Introduction

Human social networks are constructed and maintained through social interactions, as in cases of friendships, romantic ties, and business connections. Individuals build their networks by choosing (when possible) whom to interact with, varying the depth of these interactions, and connecting people in their lives in diverse and relatively stable (Saramäki et al., 2014) ways. In turn, the resulting network structures affect social interactions that take place within them. Network structures facilitate the formation of new ties, for example when individuals that have a friend in common tend to connect (Granovetter, 1973). They also constrain behaviors by modulating information access and information sharing, norm enforcement, and decision-making responsibilities and biases (Holes, 1992; Shaw, 1964; Coleman, 1988; Becker et al., 2017). Specific network configurations and compositions lead to a wide range of tangible outcomes built on social interactions, from job search success (Rajkumar et al., 2022) to substance use (Christakis and Fowler, 2008; Rosenquist et al., 2010) to political behaviors (Bond et al., 2012; Mahmoudi et al., 2024).

Social networks are formed through repeated social interactions, which unfold through mutual adjustments, (implicit) negotiation of power balance, leader-follower roles, and turn-taking patterns, among other real-time coordination mechanisms (Skewes et al., 2015; Vesper et al., 2017; Hadley et al., 2022). Experimental and observational evidence suggests that existing relationships are reflected in coordination dynamics. People who occupy more central network positions facilitate conversations more often than others when grouped with individuals from the same network (Sievers et al., 2024a), and friends and non-friends behave differently with respect to the content of conversations (Speer et al., 2024), creativity (Zhou et al., 2025), and effort contributions (Li et al., 2025a). Social ties have been also shown to modulate interpersonal physiological mechanisms during interactions, such as heartrate synchrony between closely affiliated pairs or people who know each other compared to strangers (Konvalinka et al., 2011; He et al., 2026; Andersen et al., 2026).

Recent neuroimaging studies have shown that social network properties have signatures extending beyond immediate interactions by manifesting in brain structure (Hyon et al., 2022; Kwak et al., 2018; Driver et al., 2023). For example, the degree to which undergraduate students bridge otherwise disconnected individuals as well as their centrality in the class network, are predicted by white matter connectivity in social and affective brain networks (Hyon et al., 2022). Similarly, the number of times individuals were nominated by others as discussion partners for important life topics in an whole town social network was associated with the volume of brain regions supporting social inference abilities (Kwak et al., 2018). These results suggest that individual differences in social network features have lasting neural correlates. On the one hand, the ability to navigate social situations can be used to cultivate one’s own social standing (Shin et al., 2026). On the other hand, the necessity of balancing diverse interests and points of view may gradually affect changes within brain regions linked to social expertise.

We thus hypothesized that one’s social network properties, such as e.g., network size or density, can influence how one engages in real-time interaction dynamics, potentially driven by the interplay between individual dispositions and learning from interactions within their enduring networks. If these tentative mechanisms are more broadly embodied in interaction patterns, they may also be generalizable outside of encounters with existing contacts. Supporting this idea, two neuroimaging studies found that the density of ties in personal social networks (Schmälzle et al., 2017) and the number of friends (Baek et al., 2025) were associated with activity in brain areas associated with mentalizing, social pain, and negative affect when participants played a social exclusion game with virtual partners. If present, this network–interaction link would shift our understanding of even fine-grained interpersonal behavior between strangers to be situated within social networks, and assist in team-formation and individual networking efforts, including in professional settings.

To investigate this hypothesis, we employed a simple movement task designed to capture emergence of spontaneous coordination between two people, and mimic measurable, low-level social interaction. We paired 104 strangers, and asked them to produce continuous circular movements with their right index fingers, while sitting in front of each other and seeing or not seeing the partner’s hand (a modified version of the task in Tognoli et al. (2007) and Oullier et al. (2008); Fig. 1).

**Fig. 1.**
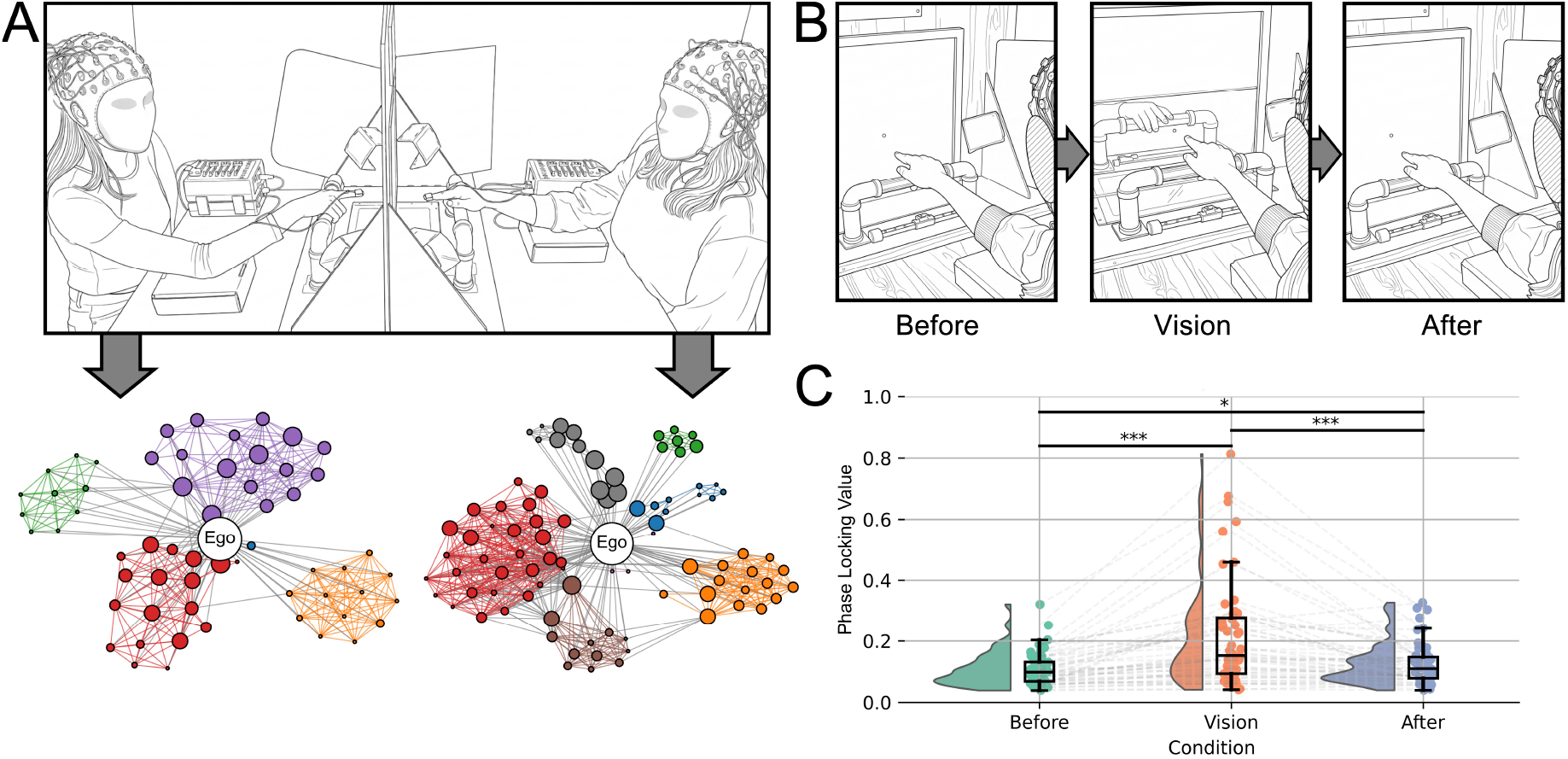
Overview of the experimental paradigm and manipulation. *(A)* The experimental setup. Participants engaged in a spontaneous coordination task while their EEG was recorded. They were separated by a screen that allowed or prohibited the view of the partner’s hand. Then each participant individually underwent a personal network mapping task and completed questionnaires. *(B)* The perspective of a participant during the three experimental conditions. *(C)* Interpersonal movement synchronization results, validating the experimental manipulation. Spontaneous movement synchronization emerged during visual contact, highlighting that participants engaged in action coordination when they could see each other’s hands (*: *P* < 0.05; ***: *P* < 0.001; based on the Wilcoxon signed-rank test with Bonferroni correction for three comparisons). The drawings of the movement task setup and participants’ perspectives in panels *A* and *B* of this figure were created using Gemini 3.1, closely following photographs of the setup.

We focused on two fundamental coordination processes with pronounced social signals: spontaneous movement synchronization and the emergence of leader-follower roles. Previous research has established that when people interact, they are likely to synchronize their movements even without instruction (Richardson et al., 2007; Tognoli et al., 2007; Hadley and Ward, 2021; Koban et al., 2019), and that being in sync has social consequences (Mogan et al., 2017), such as increased liking (Hove and Risen, 2009; Ravreby et al., 2022), cooperation (Wiltermuth and Heath, 2009; Lang et al., 2017), and prosocial behaviors (Cirelli et al., 2014). At the same time, not all spontaneous synchronization is reciprocal: one partner can show more adaptation than the other, thus taking the role of a follower, while the less adaptive partner becomes a leader (Heggli et al., 2019). Such role asymmetries feature distinct kinematic and neural signatures (Sacheli et al., 2013; Konvalinka et al., 2014; Zhou et al., 2016; Skewes et al., 2015; Li et al., 2025b). We also probed participants’ subjective assessment of the movement synchronization after they completed the movement task.

In addition to participants’ movements, we mapped their personal social networks in a separate task (inspired by Pei et al. (2022)), in which they entered their social connections across various domains (family, friends, social media contacts, etc.), then ranked them by emotional closeness and established interconnections between them. We also recorded participants’ Big Five personality traits (Johnson, 2014) as a possible mediator for network effects, since previous research has shown a relationship between network structures and personality characteristics (Centellegher et al., 2017; Rapp et al., 2019; Selfhout et al., 2010; Feiler and Kleinbaum, 2015), and personality effects on tie formation (Selfhout et al., 2010; Feiler and Kleinbaum, 2015).

To approach the neural mechanisms behind network effects on synchronization and adaptation, we recorded participants’ EEG in the hyperscanning setup. We employed multivariate temporal response functions (mTRF), which are linear filters that map the features of sensory stimuli to brain responses (Ding and Simon, 2012; Crosse et al., 2016). Such an approach was successful in describing brain responses of people dancing together (Bigand et al., 2025). In our case, mTRFs were used to predict EEG responses of interacting individuals from the movement of the partner. Additionally, we investigated inter-brain synchrony as a potential mediator between network and coordination dynamics, as the similarity of functional brain responses has been associated with relationship formation (Parkinson et al., 2018; Shen et al., 2025) and various outcomes of real-time interaction behavior (Dumas et al., 2010; Dikker et al., 2017; Fishburn et al., 2018; Astolfi et al., 2020; Nguyen et al., 2020; Atias et al., 2026).

We show that global personal network characteristics, such as community structure (the number of non-overlapping social groups), constraint (connectedness to people who already know each other (Burt, 2004)), and tie density, together with other network parameters falling on a principal component we call ’Tightness’, are associated with interpersonal motor coordination dynamics. Specifically, when leader-follower dynamics emerged, people with tighter networks (featuring fewer communities and higher network constraint) followed their partner more, while those less tight networks (more communities and less constraint) adapted less to the other, adhering to their own movement pattern. This effect was not explained by personality traits or the amount of monitoring of the partner’s movements – in fact, people with more communities and less constrained networks, who behaved more like ”leaders”, neurally encoded their partner’s movements more. This study establishes that social network properties are embodied in real-time coordination dynamics between strangers, demonstrating that awareness of the social network of the partner is not needed for the network-interaction association.

## 2 Results

### Relative Network Tightness Predicted Leader-Follower Dynamics

Fifty two pairs of strangers with the same, Danish, cultural background executed smooth circular movement with their right index fingers, with and without visual access to each other’s actions. Each dyad performed 20 experimental trials consisting of three 20-second periods: a no-vision period (“Before”), followed by a social interaction segment (“Vision”), with another no-vision segment (“After”) at the end (Fig. 1B). Participants’ movements were recorded with a 4 ms resolution to discern fine-grained behavioral dynamics.

Visual contact allowed coordinated behaviors to emerge. Significantly higher spontaneous movement synchronization, as quantified using the phase locking value (PLV) (Lachaux et al., 1999), was observed when participants could see the experimental partner’s hand (Fig. 1C). The Vision condition showed a higher median and more variable PLV (median = 0.152, IQR = 0.183) than when they had no visual feedback of each other, both in Before Vision (median = 0.098, IQR = 0.063, *P*_*Bonf*_ < 0.001) and After Vision (median = 0.11, IQR = 0.069, *P*_*Bonf*_ < 0.001) conditions. The differences between Before and After Vision conditions were also significant (*P*_*Bonf*_ = 0.011), showing that participants’ movements were significantly more synchronized after the segment with visual contact, aligned with the previously reported memory effect in a similar task (Oullier et al., 2008). The results were consistent when using the circular correlation coefficient instead of PLV (Jammalamadaka and Sengupta, 2001) (Fig. S1).

PLV in the Vision condition was positively associated with the number of trials in which participants self-reported to intentionally copy the partner (reported after completing the entire movement task; *β* = 3.35, *SE* = 0.55, z(49) = 6.1, *P* < 0.0001) and in which they felt they were in synchrony with the partner (also reported after the movement task; *β* = 2.4, *SE* = 0.52, z(49) = 4.6, *P* < 0.0001; spe- cific items and further details on the subjective measures are provided in Supplementary Information).

Next, each subject individually performed the network-mapping task, in which they listed people from different areas of their lives (family, friends, coworkers, etc.), ordered them based on emotional closeness, and specified whether the con- tacts knew each other. We derived 15 parameters from these ego networks that quantify their structure and composition (Materials and Methods). To reduce the number of variables in a simple, interpretable, and replicable way, we performed principal component analysis on 11 variables that were not an explicit subset of each other (e.g., the number of close friends is a subset of the number of friends) and were not a direct combination of others (e.g., overall network size is the sum of friends and family members) (Fig. 2A). The resulting three-component structure was replicated in an independent sample of 85 Danish and international students (Fig. S3). All principal components (PCs) featured moderate loadings but remained interpretable. The first PC captured the contrast between high density and constraint on the one side and, on the other side, the low number of communities, friends, family members, and weak ties, as well as a small circle of emotionally close contacts. We refer to this principal component as ”Tightness” of the network. The second component (”Cohesion”) was dominated by high clustering and transitivity, which indicate that a person’s contacts tend to know one another and form closed triangles. Family members and weak ties also positively loaded onto this component, suggesting that in our sample these contacts contributed to more cohesive network structures. The third component was dominated by the mean of the emotional closeness of alters (contacts) to the ego, the number of people with highest emotional closeness (close circle size), and the number of friends, which we refer to as the ”Closeness” component.

**Fig. 2.**
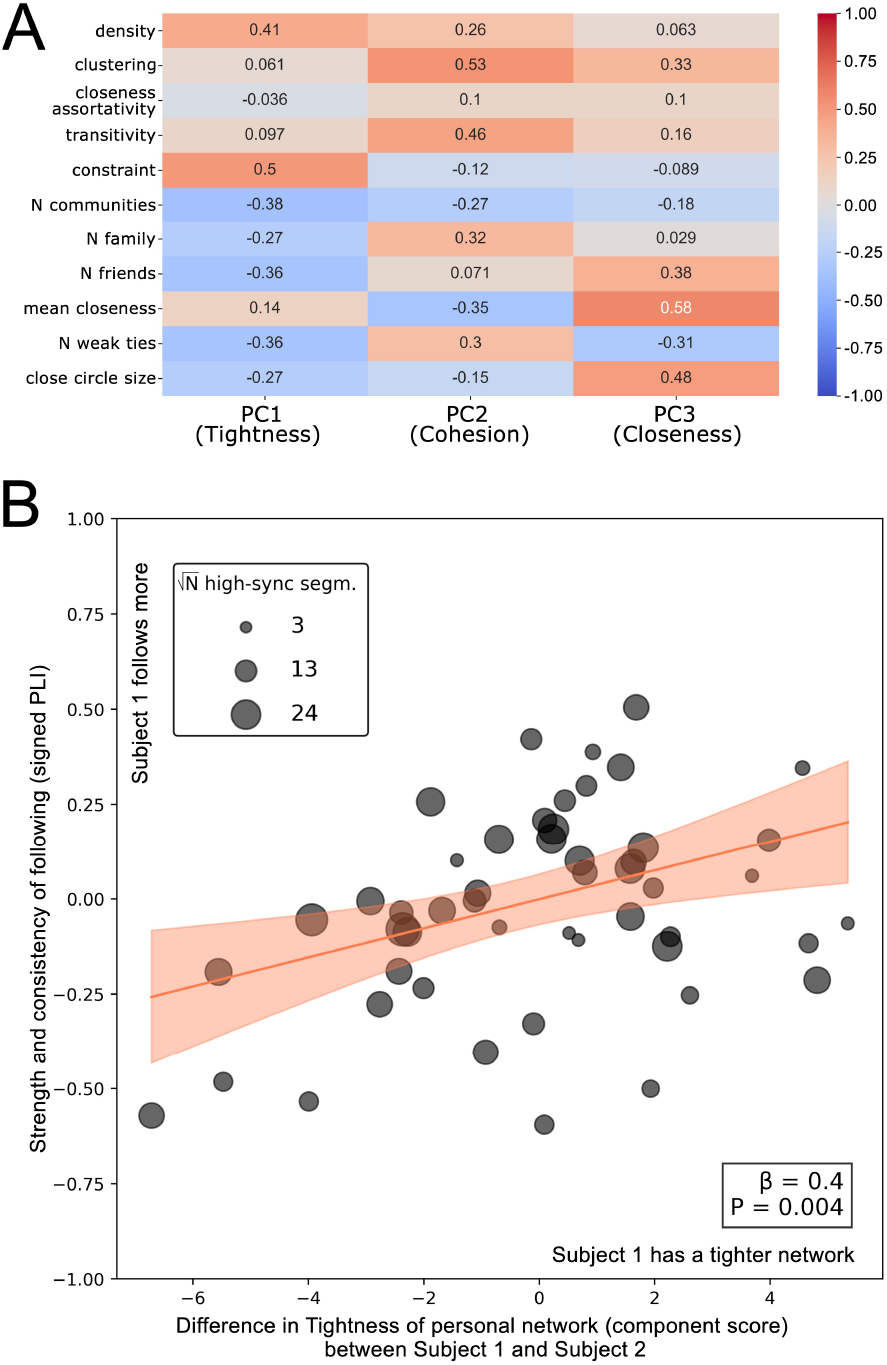
Dimensionality reduction of personal network features and the network effect on leading-following dynamics. *(A)* Loadings of principle components across ego network features. *(B)* Weighted effect of Tightness network component on leading-following behavior, controlling for Cohesion and Closeness components. Given that leading-following is an asymmetric process, we used the difference in network components between subjects in a dyad to predict the strength and consistency of relative behavioral adaptation. Participants with tighter networks than their partners (higher density and constraint but lower number of communities and friends) adapted their movements to the partner more. The -coefficient is presented here in the standardized form.

We detected behavioral leader-follower dynamics within each pair by applying the signed version of the phase lag index (PLI) (Stam et al., 2007) to the distribution of relative phase between participants’ movements during periods of high synchronization (Materials and Methods). During high synchronization, this phase-based measure quantifies the strength and consistency of the leading-following direction. Since pairs differed in the duration of high synchronization, we used weighted least-squares regression (with the square- root number of high-synchronization periods as weights). Difference in Network Tightness between interacting partners was significantly and positively associated with signed PLI (*β* _*std*_ = 0.4, *SE* = 0.13, *t*(46) = 3.001, *P* = 0.004; Fig. 2B), such that people with tighter networks than their partners were the ones who adapted more, and the higher their Tightness PC score in relation to the partner, the higher the adaptation. Differences in Cohesion and Closeness network components had no effect on adaptation (Cohesion PC: *β* _*std*_ = 0.09, *SE* = 0.12, *t*(46) = 0.754, *P* = 0.46; Closeness PC: *β* _*std*_ = 0.12, SE = 0.14, *t*(46) = 0.886, *P* = 0.38). The differences in all three PCs were entered together into a multiple regression model, in order to test whether each component predicted adaptation while controlling for the others.

Additionally, we explored whether any of the specific network parameters drove the effect of Tightness PC. We detected four trends: people with a smaller number of network communities (*β* _*std*_ = -0.49, *SE* = 0.12, *t*(49) = -3.97, *P*_*raw*_ = 0.0002) and close friends (*β* _*std*_ = -0.36, *SE* = 0.14, t(49) = -2.68, *P*_*raw*_ = 0.01), smaller close circle size (*β* _*std*_ = -0.32, *SE* = 0.14, *t*(49) = -2.32, *P*_*raw*_ = 0.025), and higher constraint (*β* _*std*_ = 0.34, *SE* = 0.13, *t*(49) = 2.57, *P*_*raw*_ = 0.013) than their partner adapted more (Fig. S4).

These leader-follower dynamics could not be explained by any differences in personality factors (Conscientiousness: *β* _*std*_ = 0.28, *SE* = 0.14, *t*(44) = 1.948, *P* = 0.058; Neuroticism: *β* _*std*_ = 0.135, *SE* = 0.16, *t*(44) = 0.855, *P* = 0.4; Extraversion: *β* _*std*_ = -0.16, *SE* = 0.15, *t*(44) = -1.09, *P* = 0.3; Openness: *β* _*std*_ = 0.07, *SE* = 0.16, *t*(44) = 0.42, *P* = 0.7; or Agreeableness: *β* _*std*_ = -0.17, *SE* = 0.16, *t*(44) = -1.09, *P* = 0.3).

Our results thus show that people with fewer network communities, smaller close circle size, and higher constraint in their personal networks (as part of a shared principal component) relative to an interacting partner adapt their movements more to the partner. Hence, personal network properties are embodied in emergent leader-follower dynamics during real-time coordination.

### People with Tighter Networks Neurally Encoded the Part- ner’s Movements Less

Next, we investigated whether a potential mechanism that could explain why participants with tighter networks adapted more, was that they more strongly monitored their partner’s movements. To capture the level of movement monitoring, we quantified how much participants neurally encoded movements of their partner. We used multivariate temporal response functions (mTRFs) to predict EEG responses based on movement stimuli (Materials and Methods). Movement stimuli entered the model jointly and consisted of three signals: self-generated movement, partner’s movement, and the instantaneous coordination between the two. The fit of mTRF models was assessed via variance partitioning: From *R*^2^ of the model containing all three behavioral components (full model), we subtracted *R*^2^ of reduced models comprising two components except the target one. To get unbiased values of *R*^2^ of full and reduced models, as well as regularization parameters (λ), we performed nested five-fold cross-validation.

The resulting models were capable of differentiating the experimental conditions (Fig. 3A). The median Δ *R*^*2*^ (across all participants) representing the unique contribution of partner-generated movements, as expected, was significantly higher during Vision than No Vision conditions (Vision-Before: *P*_*Bonf*_ < 0.001; Vision-After: *P*_*Bonf*_ < 0.0001), while the two No Vision conditions did not differ (*P*_*Bonf*_ = 0.44; *MVision* = 9 × 10^−^ 5, *MBefore* = -3.2 × 10 − 4, *MAfter* =-6.1 × 10^− 4^). The encoding of the self-generated movements lowered during Vision compared to Before visual contact (Vision-Before: *P*_*Bonf*_ < 0.0001; Vision-After: *P*_*Bonf*_ = 0.11; *M*_*V ision*_ = 9 × 10 − ^6^, *M*_*Before*_ = 0.001, *M*_*After*_ = 6.6 × 10^4^). Before and After conditions did not differ in terms of self-encoding (*P*_*Bonf*_ = 0.11). Coordination was not encoded, which is indicated by the lack of differences between experimental conditions (*P*_*Bonf*_ > 0.2) and all medians being significantly lower than zero (*P* < 0.001).

**Fig. 3.**
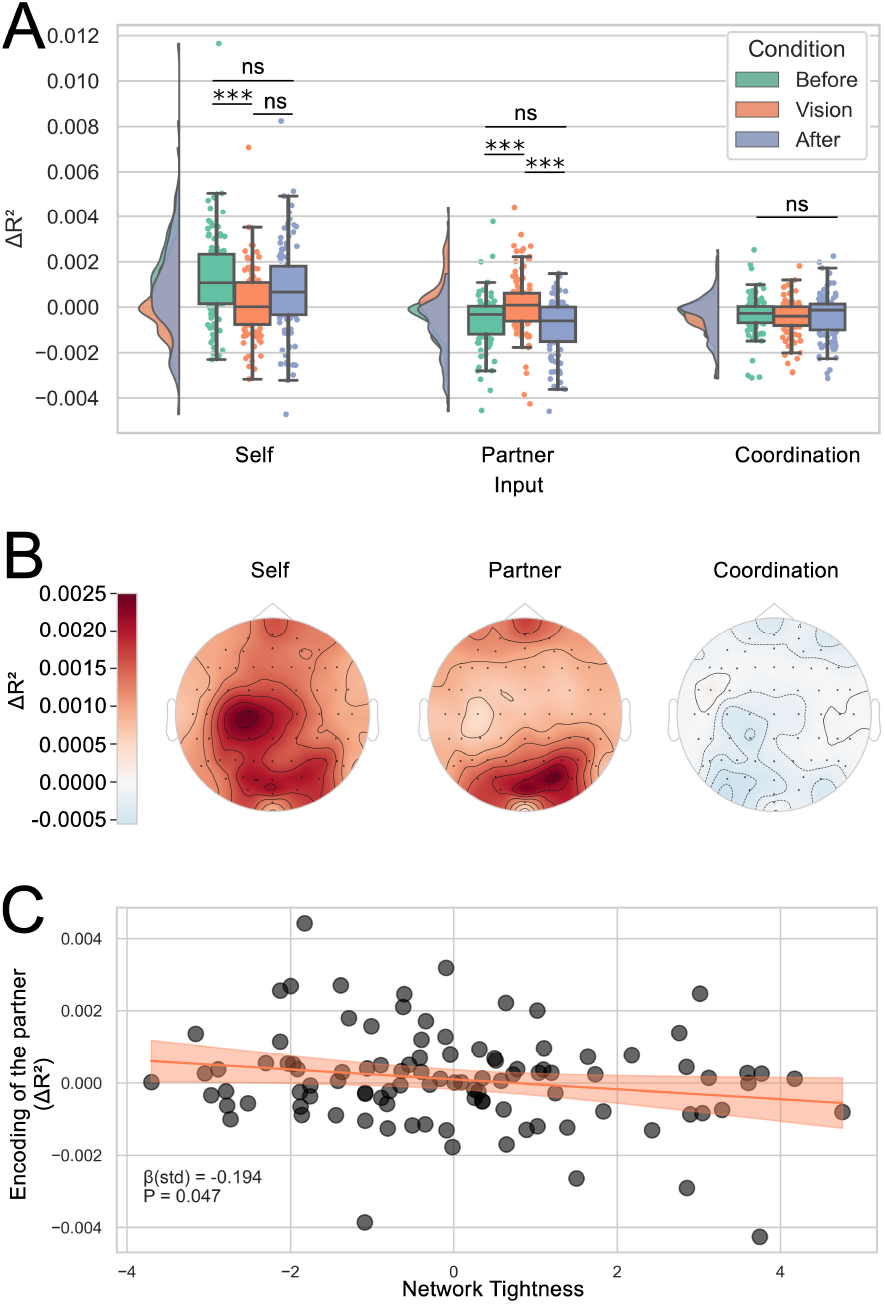
Results of fitting forward mTRFs: Participants’ EEG encoding of self-generated movement, partner-generated movement, and coordination. *(A)* Model fit for predicting EEG activity from each movement-related input type across the Before, Vision, and After conditions. As expected, partner-generated movements showed significantly higher EEG encoding when they were visually accessible. *(B)* Topographies of electrode contributions to EEG encoding during Vision per input type. Self-generated movements were largely encoded over left sensorimotor area, and electrodes in the occipital areas contributed the most to the encoding of partner-generated movements. Since coordination was poorly encoded, the topography for this input type is not informative. (C) Regression of the EEG encoding of partner- generated movements during Vision on Tightness network component, controlling for network Cohesion and Closeness. Participants with tighter networks monitored their partner less than participants with larger and sparser networks.

For visualization purposes, we reconstructed the EEG signal using the median regularization parameter coming from the folds of nested cross-validation. Then the full recordings were used to calculate R^2^ values for visualization. On average, the electrodes with the highest contribution to the encoding of self-generated movements during vision were localized over left central electrodes, corresponding to the sensory- motor area contralateral to the performing hand, with some contribution from the occipital electrodes (Fig. 3B). The encoding of the partner-generated movements was supported primarily by sensors located over occipital areas, suggesting visual encoding of the partner.

Next, we investigated whether the strength of encoding of partner-generated movements was associated with network PCs, using linear mixed models to account for the pairing of participants (Materials and Methods). Contrary to our hypothesis that higher adaptation would be associated with stronger encoding of the partner, higher partner encoding was related to a lower Tightness component score (*β*_*std*_ = -0.194, *SE* = 0.098, *z* = -1.983, *P* = 0.047; Fig. 3C), while no effects were detected for the Cohesion and Closeness PCs. This suggests that the participants with larger, sparser, and more fragmented networks encoded the partner’s movements more, even though behaviorally they adapted less.

Among personality factors, Extraversion significantly predicted the encoding of the partner (*β*_*std*_ = 0.254, *SE* = 0.11, *z* = 2.273, *P* = 0.023). Extraversion was also significantly negatively associated with the Tightness component scores (*r* = -0.34, *P* = 0.0013). However, the average causal mediation effect (ACME) of Extraversion was not established (*β*_*std*_ = -0.05, *P* = 0.15).

Self-encoding was not related to network principal components or personality factors.

Differences in self or partner encoding did not predict behavioral adaptation (Partner: *β* _*std*_ = 0.045, *SE* = 0.15, *t*(46) = 0.300, *P* = 0.8; Self: *β*_*std*_ = -0.013, *SE* = 0.15, *t*(46) = -0.084, *P* = 0.9), implying that separate mechanisms contributed to coordination dynamics and monitoring of the partner’s behavior.

### Personality Agreeableness and Personality Similarity Predicted Overall Movement Synchronization

We investigated whether network features and personality factors predicted movement synchronization during visual contact. To achieve this while accounting for the hierarchical nature of the data (each pair performed 20 trials), we used linear mixed-effects models (Materials and Methods). The average network PC scores within each pair did not predict movement synchronization (Tightness: *P* = 0.48; Cohesion: *P* = 0.27; Closeness: *P* = 0.26). In addition to the overall levels of PCs, quantified by averages, we tested whether network similarity (computed as cosine similarity across the same network features that entered PCA) predicted the amount of movement synchronization, but found only a marginal effect on PLV (*β*_*std*_ = 0.16, *SE* = 0.08, *P* = 0.059).

Instead, higher movement synchronization was associated with higher average pairwise personality Agreeableness scores, when controlling for other personality factors (*β*_*std*_ = 0.245, *SE* = 0.09, *P* = 0.005; Fig. S7). Inter- personal synchronization was also positively associated with cosine similarity across all personality facets (*β*_*std*_ = 0.19, *SE* = 0.08, *P* = 0.02), showing that partners with more similar personality traits synchronized their behavior more. Similarity in personality and network were not significantly correlated (*r* = 0.19, *P* = 0.19).

The level of synchronization was not associated with the average encoding of the partner movement (*P* = 0.69), nor by the inter-brain synchrony in phase (*P* = 0.49) or amplitude envelope correlations (*P* = 0.78) (methodological details on calculating IBS are presented in Supplementary Information).

## 3 Discussion

Our study demonstrates that properties of personal social networks manifest in moment-to-moment coordination dynamics during interactions between strangers. Individuals with relatively smaller, denser, and more constrained networks adapted their movements more strongly to their interaction partner during spontaneous motor coordination, while individuals with larger, sparser, and community-diverse networks adapted less. These effects were independent of personality differences and emerged even though participants had no prior relationship or knowledge of each other’s social networks. At the neural level, participants with large and sparse networks showed stronger neural encoding of their partner’s movements, despite behaviorally adapting less. In contrast, overall spontaneous synchronization was not predicted by social network features, but rather by personality traits and similarities between partners. Specifically, pairs with higher average personality agreeableness and personality similarity synchronized their movements more.

We propose four mutually non-exclusive mechanisms that can potentially explain the observed network effects on participants’ behavior and neural encoding: habits of engaging in interactions within a specific network structure, the ability to differentiate the self and other, sensitivity to interaction value, and seeking new connections.

### Network-specific interaction habits

The association be- tween more constrained networks and increased behavioral adaptation is consistent with the idea that different network structures place different demands on social behavior. Dense and constrained networks support interactions within limited social circles where people know each other: maintaining alignment across relationships is particularly important in such structures since trustworthiness is reciprocated and deviations from expectations are easily visible and sanctionable (Coleman, 1988). In contrast, individuals with larger and more fragmented social networks have to navigate multiple disconnected communities and social roles, which in turn allows them to behave differently with different people thus exercising greater control over their social world (Burt et al., 2002). Such conditions encourage more behavioral autonomy and may lead to reduced reliance on immediate interpersonal alignment.

### Self-other differentiation ability

The idea of behavioral and cognitive autonomy of people surrounded by large and fragmented networks is reinforced by the results of neural encoding of partner-generated movement. Although participants with such networks monitored their partners more than people with tighter networks, they were consistently less adaptive, maintaining a boundary between the self and the other. Balancing diverse points of view in daily lives may thus facilitate the ability for self-other differentiation (Koban et al., 2019) and self-concept clarity (Campbell, 1990; Campbell et al., 1996): facing a stream of new information from divergent sources, people with large and community-diverse networks may have a better understanding of how their own attitudes and behaviors are positioned in relation to other people. At the same time, a better self-other differentiation ability could be a mechanism supporting the formation of large and community-diverse networks in the first place.

### Sensitivity to interaction value

The structure of a personal network is fundamentally connected to social capital, which can be defined directly in network terms as ”resources embedded in a social structure which are accessed and/or mo- bilized in purposive actions” (Lin, 1999). Different forms of social capital can be obtained from different network structures: the Tightness network component that emerged in our study is closely related to the bonding-bridging dichotomy in the conceptualization of social capital (Putnam, 2000). While bonding capital (corresponds to small and dense networks) represents the amount of social and psychological support (Putnam, 2000; Coleman, 1988), high bridging capital (related to an ego connecting otherwise disconnected groups and alters in an unconstrained network) leads to better opportunities for ’getting ahead’ and to the higher material return of their efforts (Putnam, 2000; Burt et al., 2002; Granovetter, 1973). Thus, people with large and unconstrained networks with many disconnected communities, due to higher bridging capital, may have been less willing to invest in the limited, motor interaction that we recreated in the lab because of its low value for their real lives, which potentially demotivated them to adapt to (follow) their partner.

### Seeking new connections

People with smaller networks may have followed their partner more to include the partner as a friend into their network. Adaptive behavior, which resulted in burst of high synchronization, might have been a signal of affiliation, since movement synchronization has been shown to increase liking and cooperation (Hove and Risen, 2009; Wiltermuth and Heath, 2009; Lakens and Stel, 2011). At the same time, such behavior cannot necessarily be attributed to loneliness, because subjective feelings of loneliness and objective social isolation are typically only moderately correlated (Matthews et al., 2016; Coyle and Dugan, 2012).

While our study could not fully rule out a common cause explanation, according to which individual dispositions that influenced personal network formation also affect interaction dynamics, we found that network structure could provide information about emergent leading-following dynamics, whereas such a powerful individual characteristic as personality traits could not. This finding highlights that long-term interaction demands imposed by one’s personal network structure *can* contribute to interaction dynamics.

The results of our study point to an embodiment of social network properties in real-time interpersonal interaction behaviors. Network tightness was associated not only with motor actions per se, but also with a continuous response to another person’s actions via the sensorimotor action-perception loop. Even if a voluntary decision modulated the adaptation behavior (synchronize, resist, or ignore), this decision and its implementation to a great extent relied on the continuous sensorimotor coupling with the partner. These points align with the situatedness, action-orientedness, and time-pressure claims of embodied cognition (Wilson, 2002). Finally, the participants did not know one another prior to the experiment, were of similar age and with a similar cultural background, which were experimental design decisions aimed to limit top-down effects on interaction dynamics. During EEG application, the participants were introduced to each other and had a brief uncontrolled smalltalk to ensure that they perceived each other as human agents during the movement task. We argue that the smalltalk was unlikely to confound the network effect on leading-following because network features are not immediately observable, and a more observable characteristic, such as personality traits, was not a predictor of adaptation. Thus, participants’ behaviors were likely a response to the interaction dynamics between them.

The effect of average personality Agreeableness on the overall level of movement synchronization is only somewhat consistent with previous studies. While no effect of Agreeableness on movement synchronization was previously detected during naturalistic conversations, where movement was a supplementary modality (Arellano-Véliz et al., 2024; Tschacher et al., 2018), Agreeableness has been shown to have positive effects on more explicit measures of interaction quality (Cuperman and Ickes, 2009) and friendship formation (Harris and Vazire, 2016). The effect of higher personality similarity on higher movement synchronization can be treated as an extension of the homophily effect on human relationships, according to which similar people tend to form interpersonal connections more readily (McPherson et al., 2001; Parkinson et al., 2018), although this effect is not consistently observed (Back et al., 2011).

We should note that social network mapping by self-report, as in our study, is subject to biases, particularly node and link omission due to memory bias. We mitigated this bias by focusing only on the active network (limited to current contacts; Materials and Methods) and prompting participants with contact categories that cover major life areas, one by one. The participants were also instructed to use all possible aids (app logs, social media activity history [assisted by the experimenter]) and encouraged to focus on entering as many people as possible regardless of whether they repeated across the categories (such repetitions were later removed by the participants).

Additionally, while mTRFs are a powerful tool for matching diverse stimuli and neural responses, regressing the resulting goodness-of-fit statistics on ego-network principal components or personality factors bears additional uncertainty because, in this case, the variables both on the outcome and predictor sides are not direct measures but derivative indicators that rely on the precision of lower-level quantification steps.

Our study brings together two major developments in social neuroscience research: the move from studying isolated individuals to studying people in interaction (Redcay and Schilbach, 2019; Sebanz et al., 2006; Hari and Kujala, 2009), and the growing understanding of humans as embedded in social networks. In recent years, cognitive neuroscience has substantially contributed to this latter view by showing that network features are represented in the brain (Parkinson et al., 2017; Jensen and Parkinson, 2026) and correlate with brain structures (Hyon et al., 2022; Kwak et al., 2018) and functional brain responses (Parkinson et al., 2018; Shin et al., 2026; Schmälzle et al., 2017). Our findings extend this line of research by showing that social network properties also become embodied in low-level coordination dynamics that generalize beyond established relationships and manifest even in interactions between strangers.

## 4 Methods

### Participants

One hundred and six subjects, recruited from the general and student populations, participated in the experiment (aged between 18 and 34, mean = 23.07, SD = 3.28). The subjects were right-handed, neurotypical, with normal or corrected to normal vision. To reduce variability due to cultural factors, we recruited those who resided and were primarily raised in Denmark and spoke Danish as their mother tongue. The subjects were combined into 53 pairs of strangers, matched by age (no more than 3 years age gap between partners) and stratified by gender (15 female, 17 male, and 21 mixed). One male dyad was excluded from analysis prior to data inspection due to failure to comply with task instructions, so the final sample consisted of 52 pairs. Upon subjects’ arrival, we confirmed that they had not met before. The study was conducted according to the Declaration of Helsinki and was approved by the Scientific Ethics Committee for the Capital Region of Denmark (H-23013244). Participants were financially compensated.

### Experimental paradigm

During the movement task, subjects sat in front of each other in pairs, separated by a wall with a screen that allowed or prohibited the view of the partner’s hand via controlled opacity (Smart-film; CoolColour, Denmark). The task consisted of three conditions: Before vision (opaque screen, 20 sec), Vision (transparent screen, 20 sec), and After vision (opaque screen, 20 sec), following by a 20-sec break. The conditions were presented in the same order to control for memory effects (Oullier et al., 2008). The task consisted of 2 practice and 20 experimental trials, where each trial comprised three conditions and the break. After 10 trials, the participants had a break lasting at least 60 seconds. The participants rested their palm on a handle, and the position of their hand was adjusted so that the distance from the tip of their index finger to the screen was 6 cm (Fig. 1). The subjected were instructed to perform smooth, uninterrupted circular movement with any speed and in any direction using their index finger. It was also made explicit to them that during Vision they were allowed to synchronize with the other person and to ignore the other person, as well as to change this behavior between and within trials. To minimize EEG artifacts related to eye movements, the subjects were asked to fixate their gaze at the green circle on the separating screen, placed in the middle of expected rotation center of their partner’s finger. The trial started with an auditory signal (100 ms) and ended with an auditory cue ”Stop moving”. Similar to (Tognoli et al., 2007), participants received auditory warning cues through separate earpieces at two time points before the first 20-second interval began: 2 s (± 0.5 s, randomly distributed) and 1 s (± 0.5 s, randomly distributed). By introducing this variable delay, a random initial phase difference between participants was created, which in turn prevented shared phase priming in their movements.

### Personality questionnaire

After the movement task, the subjects were separated into different rooms (or different parts of the same room) for per- sonality questionnaire and social network assessment without ability to observe the other person’s screen. The 120-item English version of the Big Five personality questionnaire (Johnson, 2014) was presented to participants. The questions were presented in English to ensure a non-Danish speaking experimenter could answer subjects’ questions if needed, with 9 items containing idioms were accompanied by a Danish translation (Vedel et al., 2019). The order of items was pseudo-randomized per participant (plus- and minus-keyed items alternated, and factors and facets were distributed throughout the inventory).

### Ego network assessment and quantification

Participants’ active personal (ego) networks were mapped using a modified version of the Friendly Universe task (Pei et al., 2022). In this individual task, subjects were asked to enter as many people as possible whom they knew personally, where knowing personally was operationalized as knowing a person’s name, being sure they also know the subject’s name, and having interacted with them in any form within a time frame. To mitigate memory bias (Butts, 2003; Almquist, 2012), subjects entered alters in groups, presented one by one (family, friends, people from call and text logs separately, social media contacts, colleagues, face-to-face encounters, and hobby partners). The subjects were encouraged to use their smartphones to access calling, texting, and social media logs. For the family category, the time frame was not set, relying on the assumption that ties with living relatives remain at least to some extent useful throughout life. When mentioning friends, participants were instructed to list only those with whom they interacted within six months. Two-week interaction period was set for other groups. Next, subjects removed duplicating names, and ranked alters based on how emotionally close they were to the subject performing the task. The final part was to connect alters based on who knows whom personally, to the best of subject’s knowledge. To quantify the resulting ego networks, we used two sets of parameters. The first group of parameters quantified the overall network structure and included such widely-used network features as density (the ratio of present ties to the number of possible ties given network size), clustering (Watts and Strogatz, 1998), transitivity (fraction of possible triangles), constraint (Burt, 2004), closeness assortativity of alters (a tendency of similarly close alters to have a connection (Newman, 2003)), and the number of communities (more detailed definitions of the network features are privided in the Supplementary Informa- tion). Communities were identified automatically using the Louvain algorithm (Blondel et al., 2008) after removing the ego (using the Infomap algorithm (Rosvall and Bergstrom, 2008) instead of Louvain had a minimal effect on presented conclusions (Fig. S5)). The second group characterized the composition of the network by the number of family and close family members, the number of friends and close friends, the combined size of the close circle and the number of weak ties, and the mean of distribution of closeness scores in the network. We also calculated the overall size of the network and the size beyond family to specifically explore potential effects of the acquired network compared to the one the person was born into. All network features that entered PCA were log- or log(x+1)-transformed to mitigate the skew of feature distributions. The number of principal components to retain was decided based on the parallel analysis (Franklin et al., 1995).

### Movement recording and processing

Movement data were acquired using the Polhemus LIBERTY system (Polhemus, Colchester, VT). A Polhemus Micro Sensor 1.8 was attached to the tip of each participant’s right index finger. The movement data were recorded at 240 Hz sampling frequency and saved for subsequent offline analysis. The data were low-pass FIR-filtered with 10 Hz cutoff to reduce high-frequency jitter. Only the vertical dimension was used in all further analyses as it corresponds to our conceptualization of synchronization as occurring regardless of whether the movement is in the same or in different directions (clockwise or counter-clockwise). Condition triggers were sent to both movement-recording and EEG-recording equipment. Condition triggers in the movement data for one pair were reconstructed using EEG data (the triggers were placed automatically following the manual alignment of the first trigger with movement onset). The correctness of trigger placement throughout the entire recording was visually inspected.

### Quantifying movement synchronization

Phase-locking value (PLV) (Lachaux et al., 1999) was used to assess phase-based coupling (synchronization) between two signals. PLV is defined as

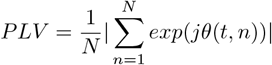

where *θ* (*t, n*) is the phase difference *ϕ*_1_(*t, n*) - *ϕ* (*t, n*).

### Detecting leading-following behavior

To detect leading-following behavior, we first identified periods of high synchronization (PLV ≥ 0.73 (Tognoli et al., 2007)) using fine-resolution sliding windows (4-seconds long with 0.5 second step). We then computed the distribution of mean relative phase by averaging relative phase values across 100 bins across all windows with synchronization above the PLV threshold. We relied only on pairs that had at least 10 windows. Two pairs did not synchronize at this threshold and were not used in the adaptation analysis. A signed version of the phase locking index (PLI) was then calculated on the resulting mean relative phase distribution, which is similar to the directed PLI (Stam and van Straaten, 2012) but ranges from -1 to 1, from lower to higher leadership, respectively. The values of signed PLI were then multiplied by -1 to reflect adaptation instead of leadership because leadership is less clearly defined in this experimental paradigm (thus, positive values mean that subject 1 adapts to subject 2). We explored the impact of window size (3—5 seconds), step size (0.5—4 seconds), PLV threshold (0.5—0.8), and the minimal number of high sync segments to enter the analysis (0—20), and conclusions remained unchanged (Table S2). PLI was used over other methods allowing for the quantification of asymmetric relationships, such as the cross-recurrence quantification analysis (Shockley et al., 2002; Konvalinka et al., 2011), because repetitive circular movement made phase-based measures of symmetric and asymmetric coupling readily-applicable and because of its simplicity and minimal researcher degrees of freedom.

### EEG recording and pre-processing

EEG data was collected in a electrically shielded room using two synchronized, daisy-chained 64-channel Biosemi ActiveTwo systems (Amsterdam, The Netherlands), arranged according to the 10–20 system and sampled at 2048 Hz. Recordings from both participants were acquired simultaneously in ActiView (v. 806) and saved for later offline analysis. Resulting EEG data were filtered to 1-40 Hz, with an addi- tional notch-filter at 50 Hz to remove residual interference from movement recoding equipment. Next, we epoched the data per condition and downsampled it to 256 Hz. After removing bad channels, we performed ICA to correct for eye blinks, horizontal eye movements, muscle artifacts, and channel noise. The data was then re-referenced to the common average, and bad channels were interpolated. Trials containing residual artifacts in the data of at least one participant in a pair were removed for the pair before further analyses (44 pairs had no such trials, 5 pairs had one trial removed, and 3 pairs had two trials removed).

### mTRF estimation

A multivariate temporal response function (mTRF) is ”a filter that describes the linear transformation of the ongoing stimulus to the ongoing neural response” (Crosse et al., 2016). Such mTRFs were estimated in windows from -250 ms to 300 ms, following Bigand et al. (2025). A separate model was fit for each person in each condition, using either all behavioral stimuli (full model) or only two of them without the target variable (reduced model). Nested cross-validation with 5 folds was performed to avoid overfitting, and a separate regularization parameter (λ; allowed range was set to 101 to 106) and a corresponding *R*^2^ as a goodness-of-fit measure were estimated at each fold. The averaged *R*^2^ across folds was used as the final goodness-of-fit measure. Finally, we used the variance partitioning approach to calculate Δ*R*^2^ for each individual stimulus type: self-generated movement, partner-generated movement, and instantaneous coordination between the two. Cosine of the instantaneous relative phase was taken as the coordination measure, and it successfully differentiated the experimental conditions in a similar fashion to PLV (Fig. S6). To fit mTRFs, we used a modified Python version of the mTRF Toolbox (Crosse et al., 2016; Bialas et al., 2023). The modifications included two aspects: parallelization of the forward model fitting to increase the speed of computation and direct calculation of *R*^2^ from model residuals. One pair, with movement triggers reconstructed using EEG triggers, did not enter the mTRF-based investigation.

### Testing differences between conditions

The overall differences in PLV between conditions were quantified using the non-parametric Wilcoxon signed-rank test and the resulting p-values were adjusted for three com-parisons using the Bonferroni correction. The Wilcoxon signed-rank test was also used to investigate whether mTRF- resulted Δ*R*^2^ per input type differed between conditions. Bonferroni correction was applied to adjust the p-values for three comparisons within each input type.

### Weighted regression

When regressing signed PLI on network principal components or personality factors to explain leading-following, weighted multiple regression was used to account for variability in the number of high synchronization segments across pairs (range = 12–603; mean = 184.9, SD = 141.8). We applied the square-root transform on the number of segments because of the window overlap (the alternative, log-transform, too aggressively equalized the weights). Weighting approach (no weights, raw number of high synchronization segments, log-transform) did not affect conclusions (Table S2). Gender did not improve model fit (*P*_*LRT*_ = 0.9), and therefore it was not included in the main model.

### Mixed-effects modeling

To estimate the effect of network components on EEG encoding of partner’s movement, we used the following linear mixed-effects model:

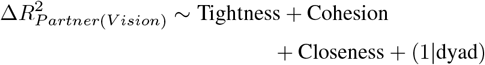

An analogous structure was used with Big Five personality factors. Based on the likelihood-ratio test, participants’ gender did not improve model fit (*P* = 0.098), and therefore it was not included in the model.

For modeling the overall level of synchronization in the Vision condition, we used linear mixed-effects beta regression models with logit link:

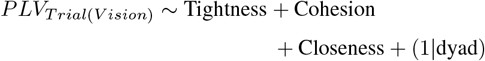

Beta regression was used because it naturally accommodated PLV values ranging from 0 to 1. The inclusion of an AR1 term to account for possible autocorrelation within a sequence of trials did not affect the results.

## Data and code availability

The behavioral, EEG, personality, and social network data, as well as the associated analysis code supporting the results of this study will be openly accessible with the journal publication of this article.

## Author contributions statement

A.D. and I.K, conceived and designed the study and experiments. A.D. conducted the experiment, collected the data, and implemented the code. A.D., I.K., and S.L. designed the analysis. A.D. and I.K. generated the first draft of the manuscript. A.D., I.K., and S.L. contributed to interpretation of the results and reviewed the manuscript.

## Declaration of Competing Interests

The authors declare no competing interests.

## Acknowledgements

This work is supported by the Villum Young Investigator grant (project no. 37525). The authors express sincere gratitude to Kiryl Vasilyeu for developing and UX-designing a standalone version of the social network task.

## Notes

### Competing Interest Statement

The authors have declared no competing interest.

