## Supplementary Information for "Personal network features are embodied in real-time social interaction dynamics"

### 1. Phase-locking value (PLV) and Circular correlation coefficient similarly differentiate experimental conditions

Phase-locking value (PLV) (Lachaux et al., 1999) and the circular correlation coefficient (Jammalamadaka and Sengupta, 2001) are used to assess phase-based coupling between two signals. Here, the circular correlation coefficient is defined as

$$\rho_c = \frac{R_{\alpha-\beta} - R_{\alpha+\beta}}{2\sqrt{Esin^2(\alpha-\mu)Esin^2(\beta-\nu)}}$$

with $R_{a\pm b} = |E(e^{i(\alpha\pm\beta)})|$ and $\mu$ and $\nu$ being the circular mean directions of phase vectors $\alpha$ and $\beta$ respectively. This implementation differs from (Burgess, 2013) but follows the reasoning and the implementation of (Jammalamadaka and Sengupta, 2001) for signals with arbitrary or uniform directions: contrary to wind and similar variables with a clear dominant direction that were fundamental for the development of the circular correlation coefficient, the phase arrays arising from the continuous circular movement do not have a well-defined direction.

The circular correlation coefficient (its absolute values, taken because anti-phase synchronization would result in negative values) is successful in differentiating the experimental conditions (Fig. S1). Higher median value was observed during the visual contact (median = 0.143) than Before Vision (median = 0.081, $P_{Bonf} < 0.0001$) and After Vision (median = 0.088, $P_{Bonf} < 0.0001$), and After Vision synchronization was significantly higher than Before Vision ($P_{Bonf} = 0.012$). The conclusions are nearly identical when compared with the outcomes of using PLV. PLV was chosen as the primary measure of synchronization due to its simplicity and popularity across studies.

### 2. Subjective assessment of movement synchronization

After the participants fully completed the movement task, they were individually asked seven questions about their own and their partner's behavior during the Vision condition:

1. In how many such rounds did you feel you were in synchrony with the other person?
2. In how many such rounds did you intentionally resist synchronizing with the other person?
3. In how many such rounds did you intentionally try to copy the other person's moves?
4. In your opinion, in how many rounds with the transparent screen did the other person resist synchronizing with you?
5. In your opinion, in how many rounds with the transparent screen did the other person try to copy your moves?
6. How much do you agree with the following statement? "I tried to synchronize with the other person, but the other person ignored me."
7. How much do you agree with the following statement? "The other person tried to synchronize with me, but I ignored the other person."

The participants could enter a number on the scale from 0 to 20, and their answers were converted into proportions. Only questions 1–3 were analyzed in this study.

The averaged within-pair proportion of trials with the subjective feeling of synchrony (question 1) and intentional copying of the partner (question 3) were significantly positively associated with PLV during Vision in a beta-regression model (feeling of synchrony: $\beta$ = 2.4, SE = 0.52, z(49) = 4.6, P $<$ 0.0001; intentional copying: $\beta$ = 3.35, SE = 0.55, z(49) = 6.1, P $<$ 0.0001; Fig. S2). Intentional resistance was not related to PLV ($\beta$ = -0.29, SE = 0.5, z(49) = -0.58, P = 0.56).

Due to a technical error, 19 pairs were presented with the stated above version of the questions, while 33 pairs had questions starting with "In what portion of such trials" (instead of "In how many such rounds"). We observed no differences in responses depending on the question formulation (Mann-Whitney U test; feeling of synchrony: P = 0.68; intentional copying: P = 0.54; intentional resistance: P = 0.1). Besides, the results presented above held in all three groups of dyads split by question formulation (pooled, first 19, after the 19th; Table S1). Hence, the responses were pooled together.

Difference in the proportion of trials in which participants reported to copy their partner was not associated with signed PLI in windows of high movement synchronization ($\beta_{std}$ = 0.157, SE = 0.12, t(48) = -1.294 , P = 0.2; Fig. S2). This discrepancy could be caused by three aspects of the task and the post-task questionnaire: (1) the participants were asked to report the number of trials with copying while leading-following was transient (signed PLI was calculated only in segments with high synchronization), (2) the intentional continuation of the sync mode might have been self-reported as a copying circle-drawing behavior (while signed PLI registered who initiated adaptation in a specific segment), and (3) at least some leading-following was driven by sensorimotor coupling occurring beneath the level of awareness.

We must also note that probing intentionality with self-report after actions occurred can provide only a limited insight into participants' cognition, as they might have ascribed intentionality to the actions post factum (without them being intentional at the moment of occurrence).

### 3. Definitions of personal network features

Fifteen features were extracted from participants' personal networks. (1) *Overall size* represented the number of contacts a person entered during network mapping. The overall size was split into (2) *the number of family members* and (3) *the size beyond family* to differentiate the part of the network a person was born into from the acquired network. Family members marked with the highest possible closeness rating (seven on the scale from one to seven) were treated as (4) *the number of close family members* feature. (5) *Friends* in this study were defined as follows: "People (NOT your relatives) with whom you currently like to spend your free time, enjoy informal social activities (such as going for lunch, drinks, films, and so on), and help each other out in a practical way. Also [...] people (NOT your relatives) with whom, in addition to spending leisure time together and helping each other out, you share personal information and provide emotional support to each other" (Burt, 1995; Parkinson et al., 2018; Sievers et al., 2024b; Spencer and Pahl, 2006). Friends with the highest possible closeness were treated as (6) *close friends*. The number of close friends, close family, and close people from other categories were combined into the (7) *close circle size* feature. The number of people with the lowest (within a given network) emotional closeness to the participant was conceptualized as the number of (8) *weak ties*. To characterize the overall tendency of a person to establish close connections, we took (9) *the mean of the closeness distribution* of the entered contacts.

Among features quantifying structural network properties, we used (10) *density*, (11) *transitivity*, (12) *clustering*, (13) *closeness assortativity*, (14) *constraint*, and (15) *the number of communities*. The ego was excluded from the network prior to computing these structural features (with an exception of constraint). *Density* was defined as the proportion of present links out of possible links given the number of nodes:

$$d = \frac{2m}{n(n-1)},$$

where $n$ is the number of nodes in the network and $m$ is the number of present edges. *Transitivity* (the fraction of triangles in the network) was calculated as:

$$T = 3\frac{Ntriangles}{Ntriads},$$

where triads are every two edges with a shared vertex and triangles are closed triads. The average *clustering* coefficient (Watts and Strogatz, 1998) for a personal network that quantified the average fraction of triangles through each node was computed as follows:

$$C = \frac{1}{n}\sum_{\nu \in G} c_\nu,$$

where $\nu$ is an individual node, $n$ is the number of nodes in network G, and node clustering $c$ defined as:

$$c_\nu = \frac{2T(\nu)}{deg(\nu)(deg(\nu)-1)},$$

with $T$ being the number of triangles through node $\nu$ and $deg$ being the degree of node $\nu$. We used the unweighted version of this and other structural measures because the participants could evaluate only the emotional closeness between themselves and their alters, while no information was collected on the alter-alter tie strength. *Assortativity*, which quantifies the extent to which similar nodes connect to each other, was calculated using the closeness attribute with the approach in (Newman, 2003) (Eq. (21)). *Constraint* is a measure that quantifies the extent to which an ego is surrounded by people who know each other (Burt, 2004). We relied on NetworkX Python package (Hagberg et al., 2007) implementation to calculate constraint on the ego (function 'constraint'), as well as for calculating density, transitivity, clustering, and assortativity. Finally, we partitioned each network into non-overlapping *communities*, which are groups of nodes with more links between them than with the rest of the network, to assess the number of social circles the ego is involved in. The partitioning was performed automatically, using the Louvain algorithm (Blondel et al., 2008). To ensure that the results are not algorithm-specific, we additionally explored whether they change if the Infomap algorithm (Rosvall and Bergstrom, 2008) is used instead of the Louvain (the outcomes of this robustness check are presented below, in section 6).

### 4. Replication of network PCA structure

Network data from 85 Danish and international students was collected using the same network mapping methodology as a part of a different study in our lab. The principal component structure resulting from the main data set and from this additional data closely resembled each other (Fig. S3), pointing to its robustness.

### 5. Individual trends in the relations between network features and leading-following

The supplementary plots showing the trends in the relations between network features and adaptation are presented in Fig. S4.

### 6. Algorithms for network community detection: Infomap leads to similar conclusions as Louvains

Changing the community detection algorithm from Louvain to Infomap has lead to nearly identical main results (Fig. S5). The effect of cosine similarity across network features (network similarity) on the overall level of trial synchronization has crossed the significance threshold with Infomap ($\beta$ = 0.165, SE = 0.08, $P$ = 0.044).

### 7. Robustness examination of the relationship between Tightness PC and leading-following

Table S2 summarizes the results of robustness checks of:

- the weighting approach in the weighted least squares regression,
- step size of the sliding window for detecting high-synchronization segments,
- window size of that sliding window,
- PLV threshold for considering a segment a high-synchronization or not,
- minimal number of high-synchronization segments for a pair to enter the analysis.

The outputs are supplemented with the number of retained pairs resulting from a specific parameter value.

### 8. Instantaneous coordination measure for mTRF

Cosine of relative phase was used as an instantaneous coordination measure for mTRF fitting. It successfully differentiated Vision from No Vision conditions (Fig. S6).

### 9. Personality effects on movement synchronization

Personality Agreeableness and overall personality similarity were positively associated with movement synchronization (Fig. S7).

### 10. Results of inter-brain synchrony

To ensure the interpretability of inter-brain synchrony (IBS), we focused on homologous regions between two brains of the interacting partners. We investigated the association between movement synchronization and IBS in:

1. the alpha band (8-13 Hz) over the occipital area (Oz, O1, and O2 electrodes) to probe the involvement of visual attention (Jensen and Hanslmayr, 2020; Klimesch, 2012),
2. the left and right mu band (8-13 Hz; electrodes C3 and C4 respectively) due the potential involvement of the activity in this band in the mirroring mechanism (Muthukumaraswamy et al., 2004; Fox et al., 2016),
3. the beta band (14-30 Hz) over the right and left sensorimotor areas (electrodes C2, C4, CP2, CP4 and C1, C3, CP1, CP3, respectively) as the activity in this band is involved in motor control (Engel and Fries, 2010),
4. the mid-frontal theta band (4-7 Hz; electrodes Fpz, AFz, Fz, and FCz) as it is involved in cognitive control (Cavanagh and Frank, 2014).

We quantified brain-to-brain coupling in these areas and frequency bands both in phase, using PLV (Dumas et al., 2010), and as the amplitude envelope correlation (Zamm et al., 2018). For the quantification, we removed the initial two seconds of each trial to avoid spurious coupling caused solely by the change in the mode of the separating screen. Then, the trials (now 18-sec long) were split into 6-second segments both in EEG and movement (Zimmermann et al., 2024). To test the association between IBS and movement synchronization, we fit mixed-effects models of the following structure:

$$IBS_{segment} \sim PLV_{segment} * FROI + diag(1 + FROI|pair : trial),$$

where $FROI$ stands for the frequency-region of interest. The "diag" command in the formula constrains the random effect structure to be fit with zero correlation between the random slope and random intercept, the necessity of which was dictated by convergence issues of the model. A term accounting for a possible autocorrelation between segments (because each three segments came from the same trial) was not used due to convergence issues. We used the beta regression with logit link for the phase-based IBS, due to IBS values being strictly positive and not greater than 1; the gaussian model was used for the amplitude-based IBS. Neither phase-based IBS (P=0.49) nor amplitude-based IBS (P=0.78) were predicted by movement synchrony in the interaction with the frequency-region of interest.

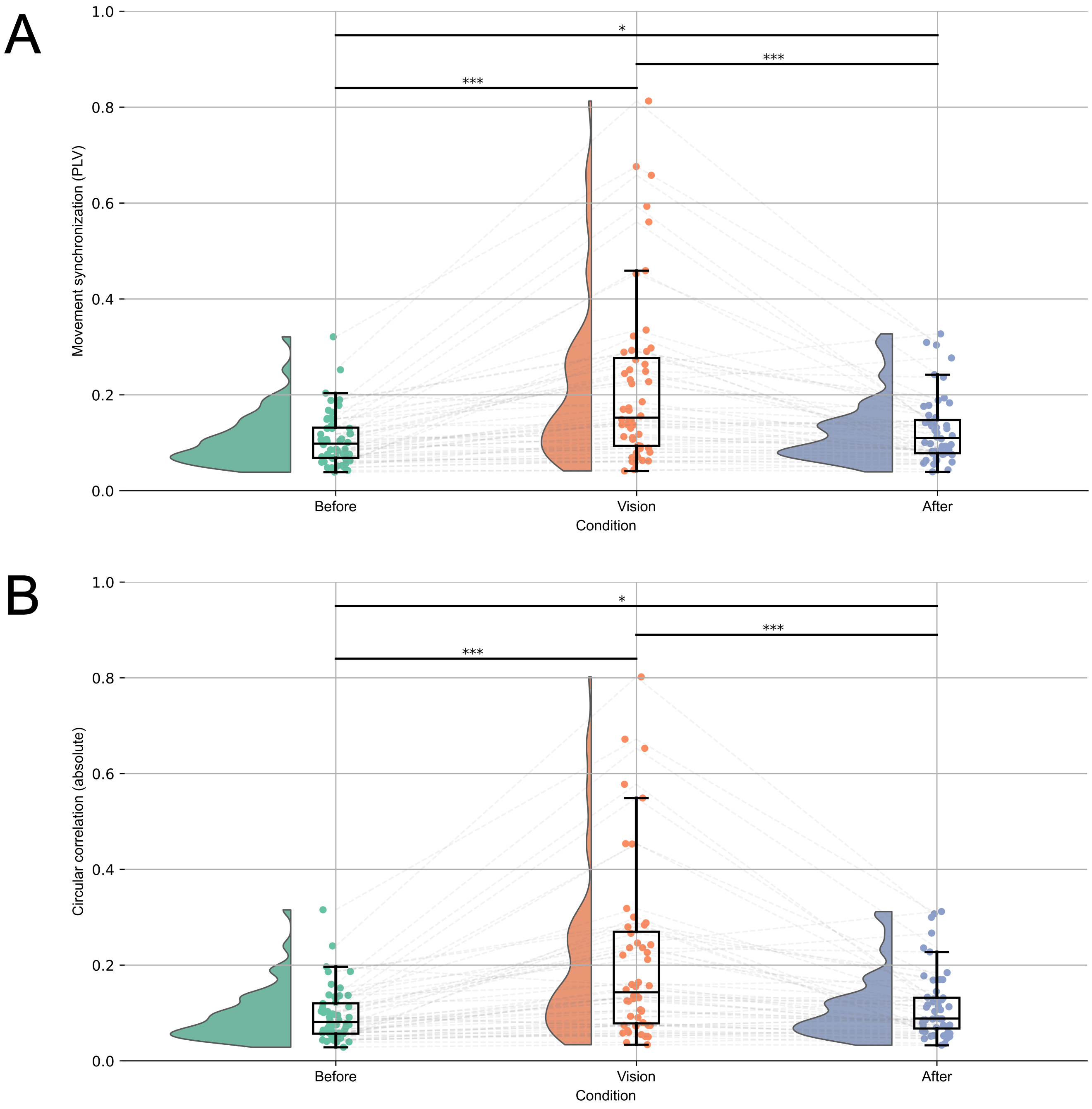


**Fig. S1**. Comparison of performance of two measures for phase-based coupling of movement signals in the current experimental setup. *(A)* Phase-locking value and *(B)* Circular correlation coefficient produce nearly identical conclusions.

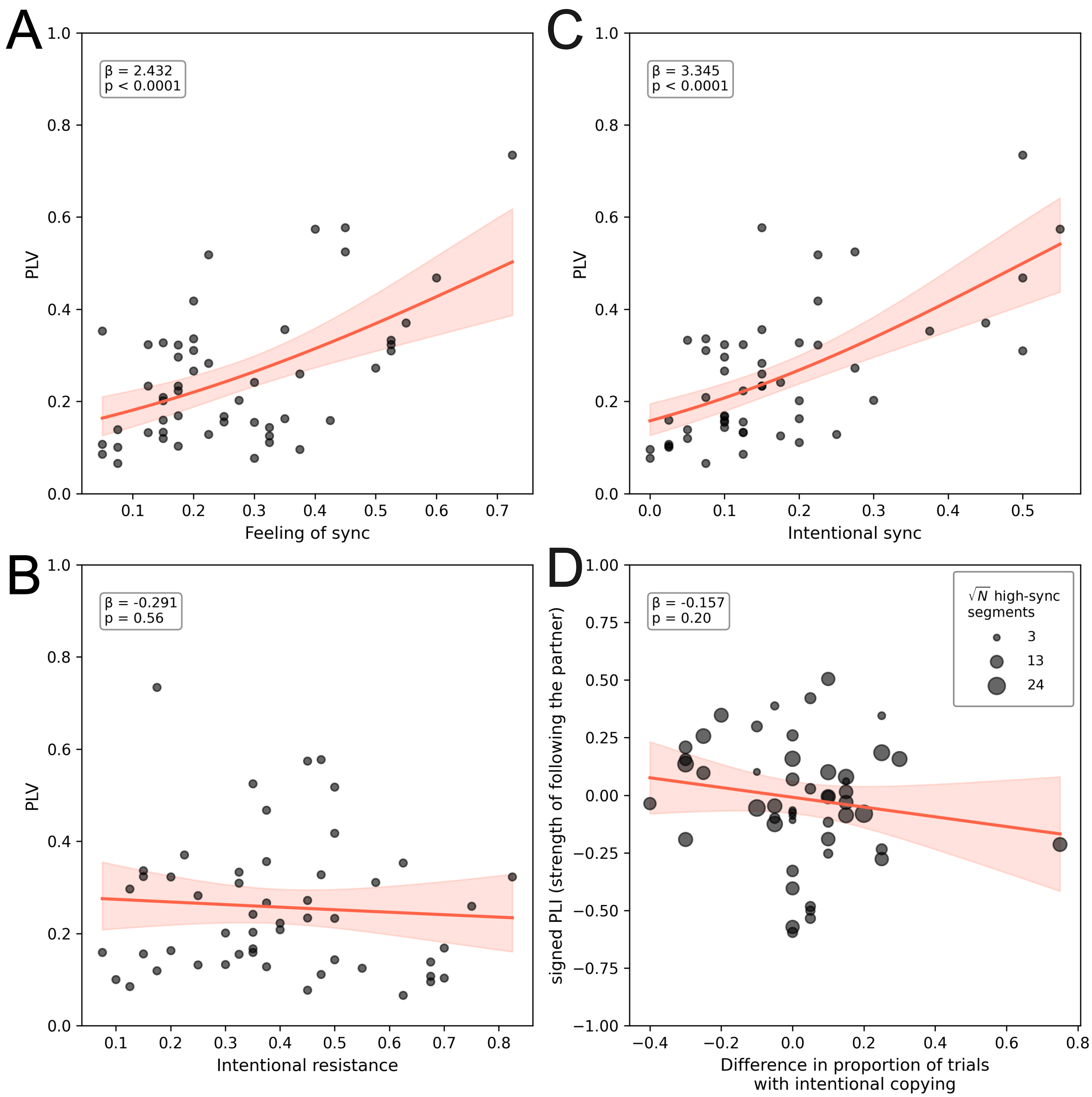


**Fig. S2**. Relations between the self-reported assessment of synchronization behaviors and the measures of symmetric *(A, B, C)* and asymmetric *(D)* coupling. The dyad-averaged proportion of trials where the participants felt in synchrony *(A)* and where they reported to intentionally copy the partner's movements *(C)* was significantly associated with PLV, highlighting the internal validity of the measure. Self-reported intentional resistance to synchronize *(B)* was not related to PLV. Difference in the number of trials where the subjects reported they copied the partner's movements was not associated with leading-following behavior *(D)*.

**Table S1**. Association between PLV and subjective measures split by question formulation

| | Pooled | | | | First 19 | | | | After 19 | | | |
|---|---|---|---|---|---|---|---|---|---|---|---|---|
| | $\beta$ | SE | Z | p | $\beta$ | SE | Z | p | $\beta$ | SE | Z | p |
| **Feeling of sync** | 2.432 | 0.525 | 4.634 | $< .0001$ | 2.764 | 0.647 | 4.272 | $< .0001$ | 2.217 | 0.739 | 3.000 | 0.003 |
| **Intentional sync** | 3.345 | 0.547 | 6.120 | $< .0001$ | 3.057 | 0.672 | 4.549 | $< .0001$ | 3.578 | 0.869 | 4.119 | $< .0001$ |
| **Intentional resistance** | -0.291 | 0.505 | -0.578 | 0.564 | 0.058 | 1.021 | 0.057 | 0.955 | -0.222 | 0.596 | -0.372 | 0.710 |

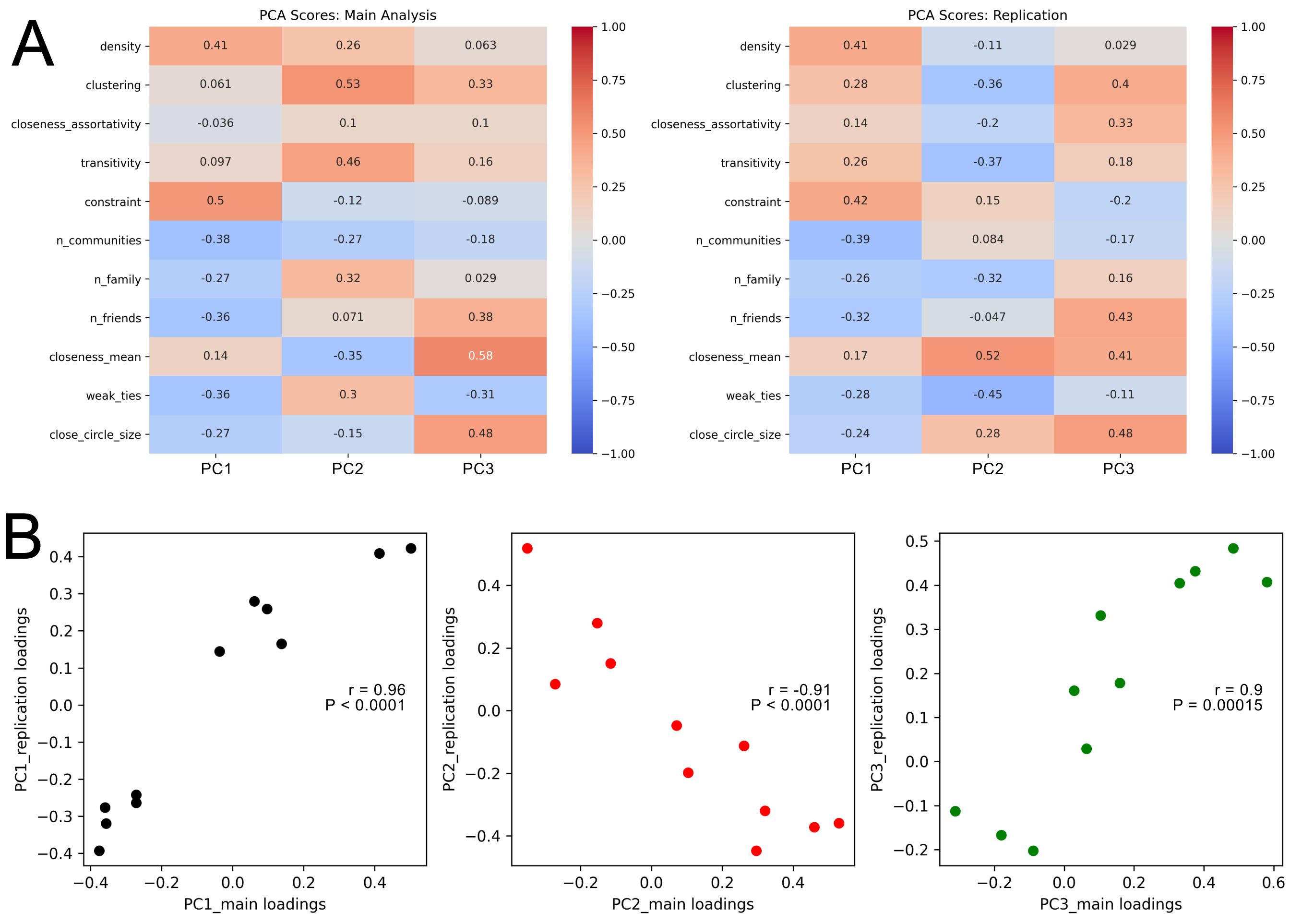


**Fig. S3**. Replication of the principal component structure in an additional data set of 85 Danish and international students. *(A)* Loadings of the principal components. *(B)* Correlations between the main and replication principal component loadings. The inverse direction of the second principal component in the replication data set supports replication despite the opposite sign because PC signs in the PCA are arbitrary.

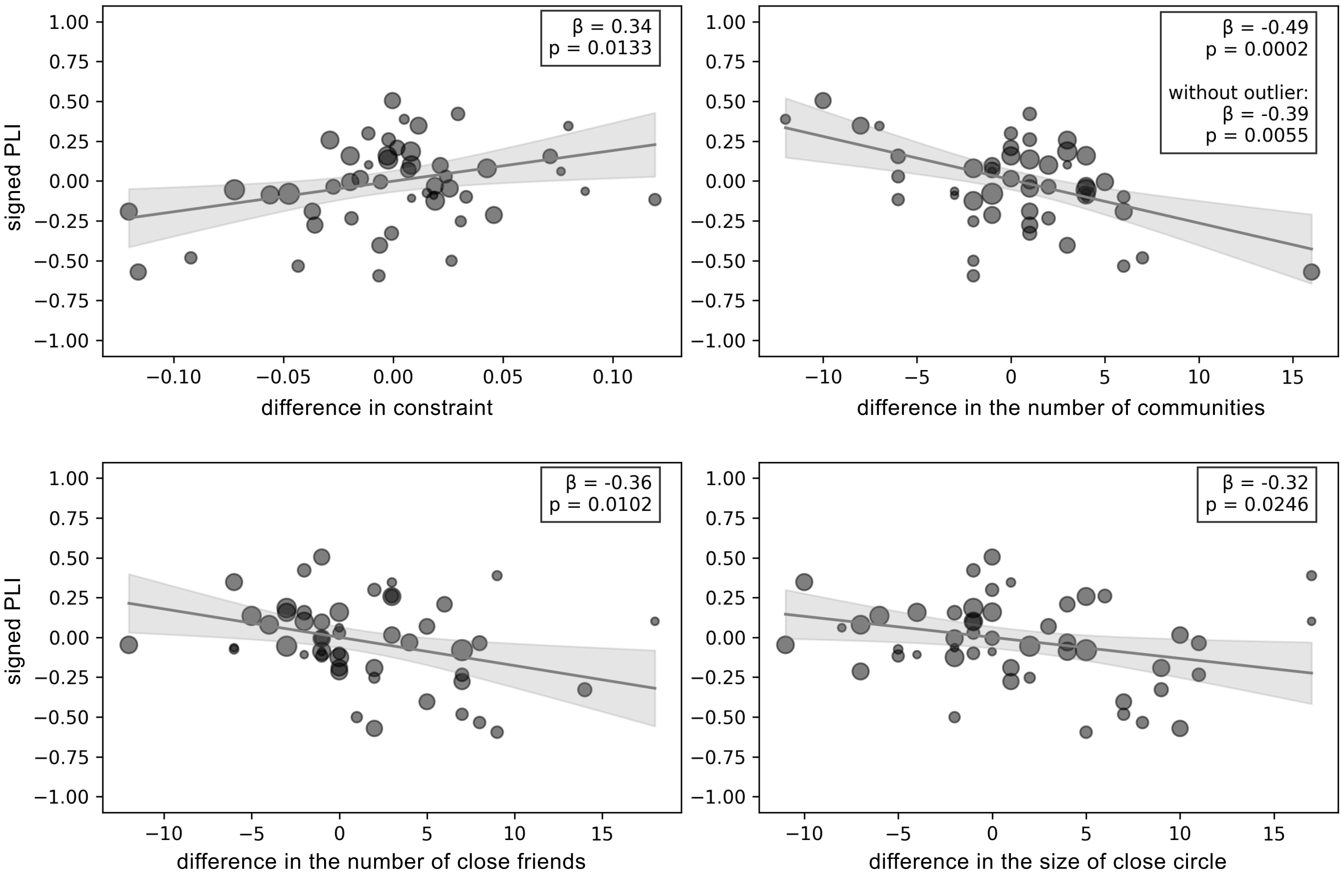


**Fig. S4**. Trends in the association between leading-following strength (signed PLI) and the difference between interacting partners in network features. The other eleven network features showed no effect on leading-following. The coefficients are standardized, and the p-values are presented here without an adjustment for multiple comparisons.

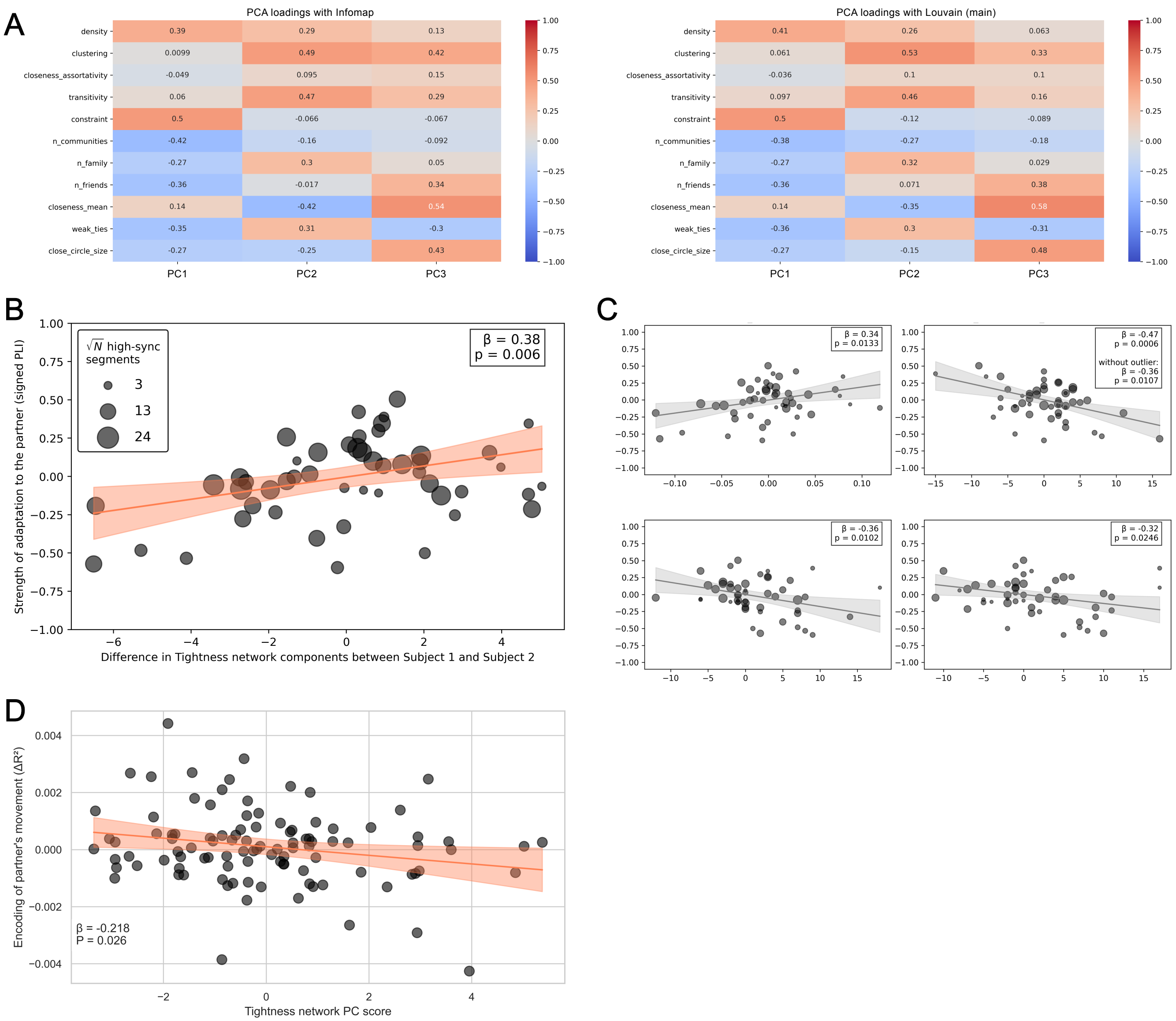


**Fig. S5**. The changes in key results due to swapping the Louvain community detection algorithm for the Infomap algorithm. *(A)* PCA loadings remain very similar. *(B)* The effect of network Tightness PC (PC1) changed minimally. *(C)* The number of communities as a pre-PCA feature still has the same trend. *(D)* The effect on network Tightness on partner encoding (mTRF) remained unchanged.

**Table S2**. Post-hoc robustness examination of the association between Tightness PC and leading-following (signed PLI)

| Weighting approach | $\beta_{std}$ | SE | t | P | N observations |
|---|---|---|---|---|---|
| square root of N high-sync segments (main) | 0.396 | 0.132 | 3.001 | 0.004 | 50 |
| no weights | 0.388 | 0.136 | 2.849 | 0.007 | 50 |
| raw N high-sync segments | 0.392 | 0.125 | 3.135 | 0.003 | 50 |
| log N high-sync segments | 0.397 | 0.136 | 2.925 | 0.005 | 50 |
| Step size (sec) | $\beta_{std}$ | SE | t | P | N observations |
| 0.5 (main) | 0.396 | 0.132 | 3.001 | 0.004 | 50 |
| 1 | 0.385 | 0.134 | 2.880 | 0.006 | 45 |
| 2 | 0.335 | 0.145 | 2.318 | 0.026 | 42 |
| 3 | 0.407 | 0.141 | 2.889 | 0.006 | 40 |
| 4 | 0.441 | 0.149 | 2.948 | 0.006 | 37 |
| Window size (sec) | $\beta_{std}$ | SE | t | P | N observations |
| 3 | 0.366 | 0.128 | 2.860 | 0.006 | 52 |
| 4 (main) | 0.396 | 0.132 | 3.001 | 0.004 | 50 |
| 5 | 0.372 | 0.131 | 2.841 | 0.007 | 45 |
| PLV threshold | $\beta_{std}$ | SE | t | P | N observations |
| 0.5 | 0.356 | 0.125 | 2.842 | 0.007 | 52 |
| 0.6 | 0.372 | 0.124 | 3.003 | 0.004 | 51 |
| 0.7 | 0.397 | 0.123 | 3.231 | 0.002 | 51 |
| 0.73 (main) | 0.392 | 0.125 | 3.135 | 0.003 | 50 |
| 0.8 | 0.348 | 0.121 | 2.873 | 0.006 | 45 |
| N high-sync segments threshold | $\beta_{std}$ | SE | t | P | N observations |
| 0 | 0.394 | 0.128 | 3.075 | 0.003 | 52 |
| 5 | 0.398 | 0.131 | 3.042 | 0.004 | 51 |
| 10 (main) | 0.396 | 0.132 | 3.001 | 0.004 | 50 |
| 15 | 0.390 | 0.133 | 2.931 | 0.006 | 45 |
| 20 | 0.390 | 0.133 | 2.931 | 0.006 | 45 |

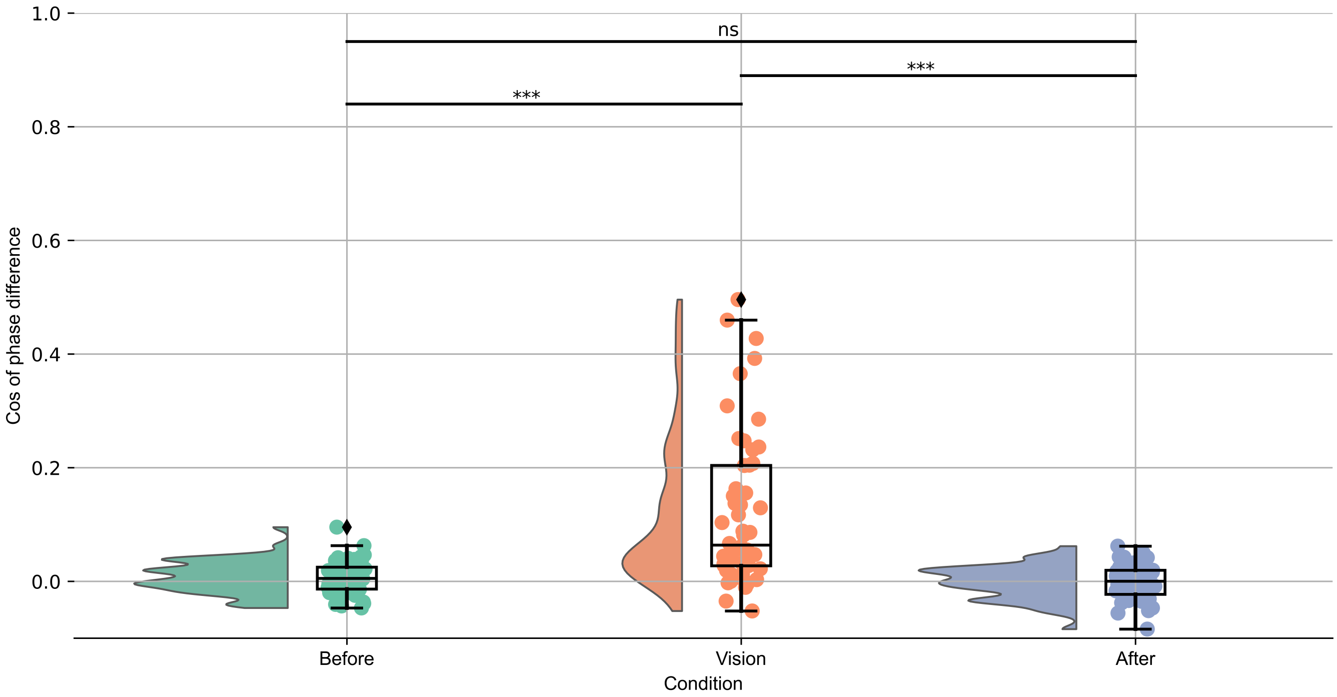


**Fig. S6**. Cosine of phase difference as a coordination measure. Cosine of phase difference, which served as an instantaneous measure of coordination for mTRF fitting, was capable of differentiating Vision from No Vision conditions (averaged per pair to be comparable with PLV results). P-values are based on Wilcoxon signed-rank test with Bonferroni correction (***: $P < 0.001$).

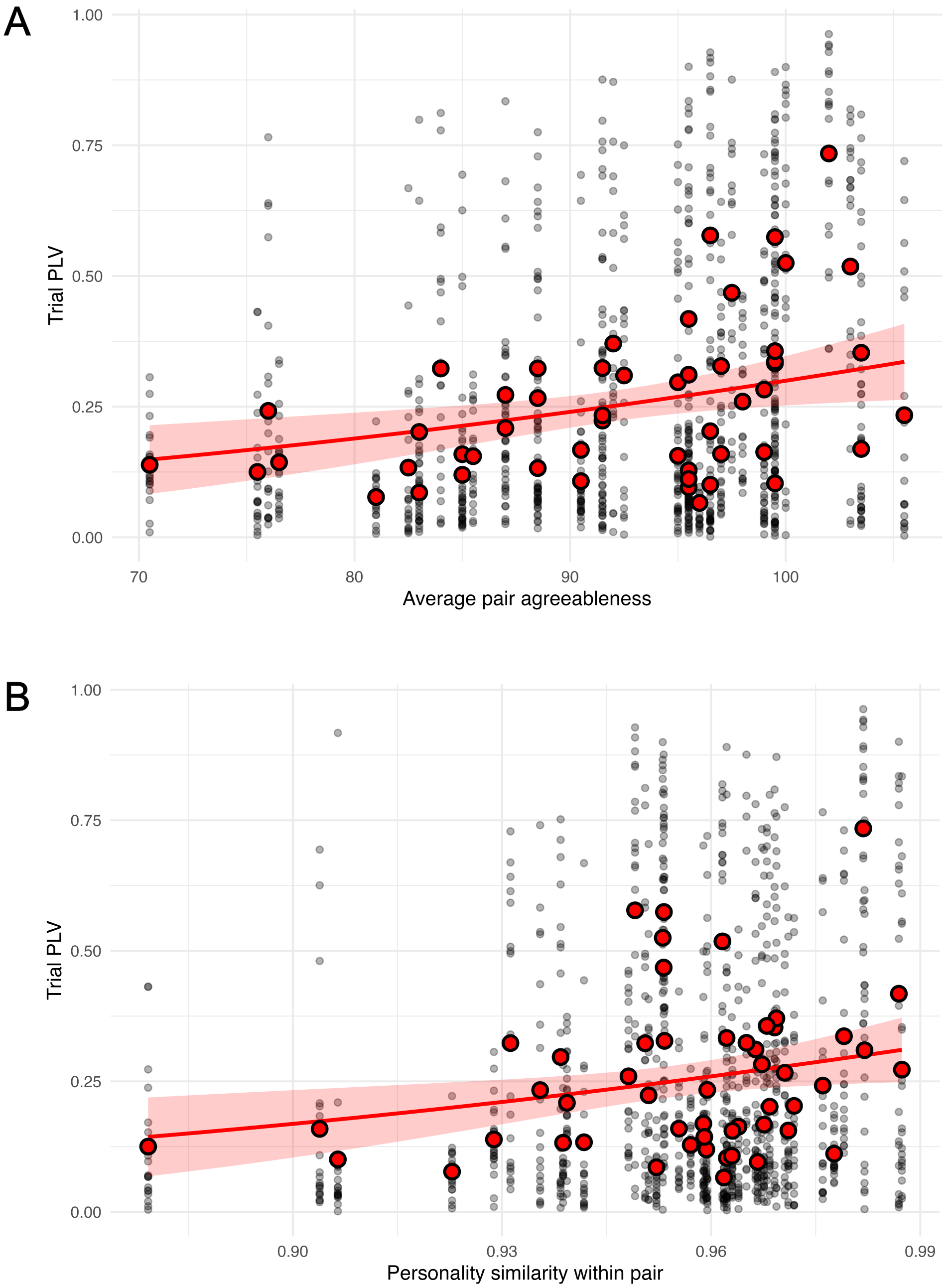


**Fig. S7**. Personality effects on movement synchronization (PLV). *(A)* The effect of the averaged within pairs Agreeableness scores on PLV. Black circles represent the data per experimental trial (20 data points per pair). Red circles represent averaged PLV per pair across the entire experiment. The red line depicts the fit of the beta mixed-effects regression model. *(B)* The effect of personality similarity (cosine similarity across 30 Big Five facets within pair) on movement synchronization. The same notation applies as in *(A)*. The removal of outliers (z-score above $|3|$) on the personality similarity axis as a robustness check did not affect the conclusions (without the outliers: $\beta_{std} = 0.169, SE = 0.08, z = 2.054, P = 0.04$).
